# Mechanism of anterior cruciate ligament loading during dynamic motor tasks

**DOI:** 10.1101/2020.03.15.992370

**Authors:** Azadeh Nasseri, David G Lloyd, Adam L Bryant, Jonathon Headrick, Timothy Sayer, David J Saxby

## Abstract

This study determined anterior cruciate ligament (ACL) force and its contributors during a standardized drop-land-lateral jump task using a validated computational model. Healthy females (n=24) who were recreationally active performed drop-land-lateral jump and straight run tasks. Three-dimensional whole-body kinematics, ground reaction forces, and muscle activation patterns from eight lower limb muscles were collected concurrently during both tasks, but only the jump was analyzed computationally, with the run included for model calibration. External biomechanics, muscle-tendon unit kinematics, and muscle activation patterns were used to model lower limb muscle and ACL forces. Peak ACL force (2.3±0.5 BW) was observed at 13% of the stance phase during the drop-land-lateral jump task. The ACL force was primarily developed through the sagittal plane, and muscle was the dominant source of ACL loading. The gastrocnemii and quadriceps were main ACL antagonists (i.e., loaders), while hamstrings were the main ACL agonists (i.e., supporters).

## Introduction

Rupture of the anterior cruciate ligament (ACL) is one of the most common and debilitating knee injuries. In Australia, ACL ruptures result in an annual ACL reconstruction (ACLR) prevalence of 77.4 per 100 000 individuals, which is the highest in the world, and has been increasing in adolescents and young adults over the past decades [1]. The majority of ACL ruptures do not involve direct collision, but occur during landing, cutting, and pivoting tasks common to sports such as soccer, basketball, and netball [2]. Moreover, ACL ruptures are 3.5-4 times more frequent in female compared to male athletes [3]. Multi-planar knee loading is associated with elevated ACL loading, suggested from laboratory biomechanical studies [4], *in vivo* measurement [5] and cadaveric experiments [6]. However, rarely have biomechanical studies examined ACL loading directly or included the role of the individual’s muscle activation patterns in stabilising the knee. Consequently, the precise mechanisms behind ACL loading during dynamic activities are unclear, thus depriving injury prevention and rehabilitation programs of personalized targets mechanistically linked to *in vivo* ACL loading.

There are few studies reporting direct measurement of ACL loading during dynamic tasks such as jumping and landing [7, 8] and similarly few using indirect image-based assessments [9]. Computational models, if validated, enable non-invasive assessment of ACL loading during dynamic tasks for large cohorts and, potentially, function of muscle in loading the ACL. Previous computational models were developed based on small sample size data [10], assessed simplistic knee loading conditions [11], and/or did not predict experimentally measured ACL loading [12]. Recently, a phenomenological model of ACL loading during dynamic tasks was developed, validated, and applied to a standardized land and jump task [13]. Findings demonstrated ACL force was primarily generated through the sagittal plane mechanism, with muscle being the main contributor due to their lines of action. However, this prior study [13] was limited to a small sample of individuals and did not thoroughly explore the contributors to ACL force, leaving the role of specific muscles in loading and/or supporting the ACL unanswered.

Considerable research focus has been devoted to the role of the quadriceps and hamstrings in loading and/or supporting the ACL [14] as their anatomical relationships with the knee has made them logical priorities for investigation. However, comparatively little research has focused on the role of the gastrocnemii on ACL loading during dynamic tasks. Cadaveric studies [15] indicate the gastrocnemii load the ACL in non-weight bearing conditions, which is consistent with modelling studies [16]. Finite element modelling also identify the gastrocnemii as ACL antagonists (i.e., loaders) under simplistic motion and boundary conditions such as isometric contraction at different force levels or knee angles at during specific instances of stance [17]. However, no previous study has examined ACL force during dynamic tasks with a validated model that accounts for the individual’s muscle activation patterns (i.e., subject-specific activations).

The purpose of this study was to determine ACL forces during a standardized drop-land-lateral jump task in females using a validated computational model that includes subject-specific muscle activation patterns, second, to determine the contributions of muscle to ACL force, as muscle recruitment could theoretically be modified by specialized training. We hypothesized: based on previous studies of *in vivo* landing mechanics [13, 18] and cadaveric experiments [19], the ACL will receive the majority of its loading via the sagittal plane mechanism; based on cadaveric experiments [19], we expect the quadriceps to be the primary ACL loader (i.e., antagonists) and the hamstrings to be the main supporters (i.e., agonists).

## Methods

This study was approved by the University of Melbourne Human Research Ethics Committee (#1442604) and data acquisition was conducted at the Centre for Health, Exercise & Sports Medicine, University of Melbourne, Australia. Twenty-four healthy females (age: 19.95±4.07 years; mass 59.79±9.34 kg; height 1.65±0.06 m; body mass index 21.9±3.42 kg. m^-2^) volunteered to participate in this study. All participants were recreationally active and had no history of lower limb injury, knee pain, or previous ACL or meniscal injuries. All participants were classified as late/post-pubertal based on Tanner’s pubertal classification system (i.e., Tanner stages IV and V). Participants, or their parents/guardians for those <18 years of age, provided their informed consent before data collection.

Each participant attended a laboratory-based testing session, wherein three trials of a standardized drop-land-lateral jump from a box, with a height of 30% of lower limb length, and three trials of running with their natural running style (speed 2.8-3.2 m. s^-1^). The running data were used in subsequent model calibration (described later). Box height was normalized to each participant’s leg length to impose a similar task demand across individuals of different stature. For the drop-land-lateral jump, participants began by standing on their non-dominant leg at the centre of the box 10 cm from the force plate and folding their hands across their chest. Participants were instructed to drop down and land on their non-dominant leg on the marked target at the centre of the force plate, and immediately perform a 90° lateral jump landing on their contralateral leg on the marked target which was at a distance of 150% lower limb length from the centre of the force plate. Participants were asked to perform the drop-land-lateral jump task consistent with the instructions, but as quickly as possible. The selection of this task was based on its similarity to the landing manoeuvre in sports and given that majority of ACL ruptures occur during a single leg stance [20].

During these tasks, body motion, ground reaction forces (GRF), and EMG were synchronously and concurrently acquired. Body motion was acquired using a 12-camera motion capture system (Vicon Motion Systems, Oxford, UK) sampling at 120 Hz, body-ground interaction was acquired using ground embedded force platforms (AMTI, Massachusetts, USA) sampling at 2400 Hz, and muscle activations were acquired using a wireless surface EMG system (Noraxon, Arizona, USA) sampling at 2400 Hz. Each participant was outfitted with retroreflective markers mounted on specific anatomical landmarks, described in [21]. Surface EMG electrodes were applied overlying the eight major lower limb muscles (i.e., rectus femoris, vastus lateralis, vastus medialis, tibialis anterior, lateral gastrocnemius, medial gastrocnemius, lateral hamstrings, and medial hamstrings) of the non-dominant leg consistent with the guidelines from Surface Electromyography for the Non-Invasive Assessment of Muscles (http://www.seniam.org/).

All data were filtered using second-order, zero-lag, Butterworth filters. Body markers and GRF were low pass filtered with 6 Hz cut-off frequency. Raw EMG was band-pass filtered (30-300 Hz), full-wave rectified, and then low-pass filtered with a cut-off frequency of 6 Hz to produce linear envelopes. Each linear envelope was subsequently normalized to its maximum envelope value identified from all available trials. The overall processing flow involved in using laboratory human motion data to estimate ACL force is displayed in a block diagram (Figure 1). We will begin our explanation with the use of OpenSim [22] version 3.3 to model the external biomechanics during the drop-land-lateral jump task. A full-body generic musculoskeletal model was used, consisting of 37 degrees-of-freedom (DOF) and 80 muscle tendon unit (MTU) actuators [23]. This generic model was modified by first adding dummy tibias of negligible mass and associated universal joints without changing the original knee mobility of flexion/extension with other motions (i.e., abduction/adduction rotation, internal/external rotation, superior-inferior translation, and anterior/posterior translation) prescribed as functions of knee flexion, and second expanding the generic model’s ankle and hip joints to 6 DOF, with the newly expanded DOF having zero mobility space. As a result, the ankle had plantar/dorsi-flexion rotational mobility and the hip had flexion/extension, adduction/abduction, and internal/external rotational mobilities.

**Figure 1.**
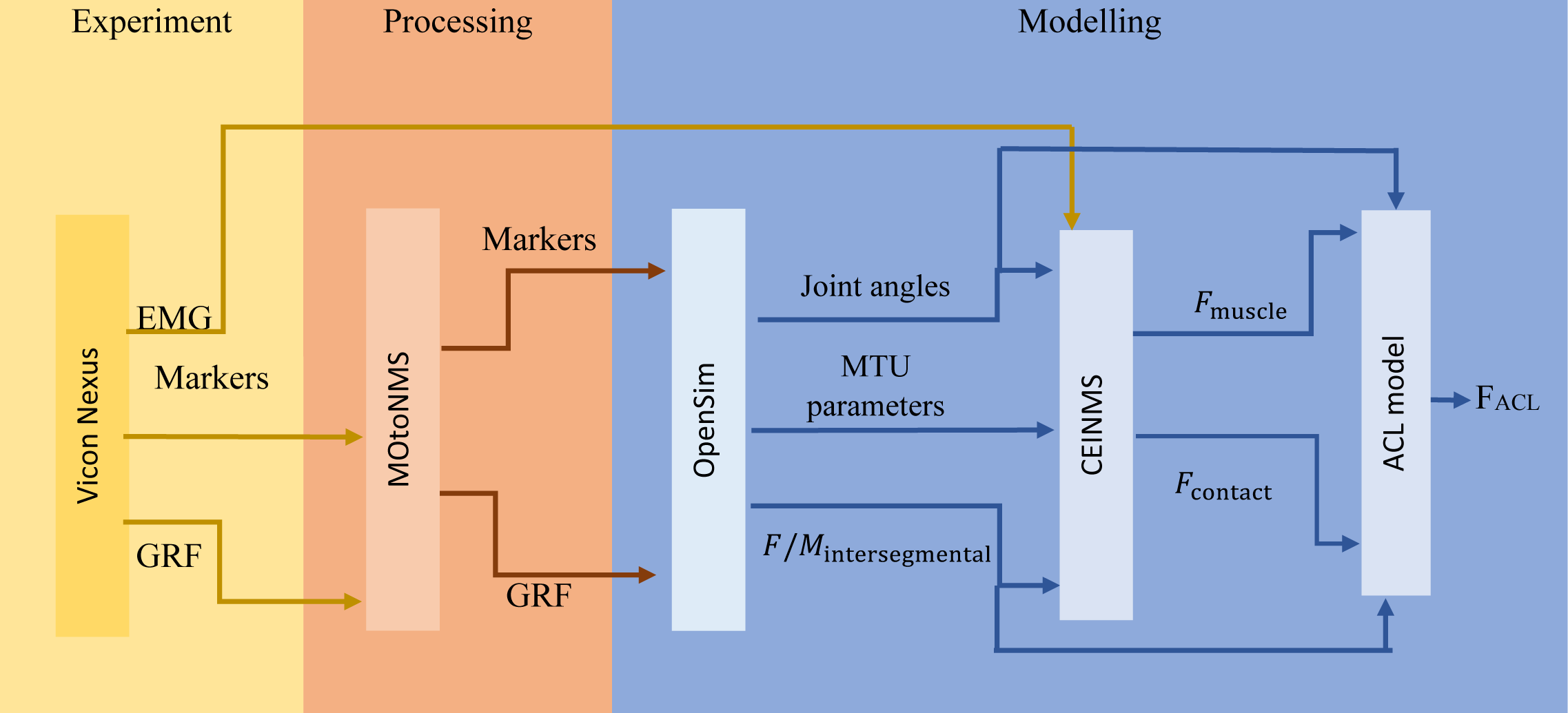
Schematic of ACL force calculation workflow. It is composed of three main parts: experimental acquisition of movement data (yellow), data processing (orange), EMG-driven musculoskeletal modelling and ACL force calculation (blue). The box blocks represent the processing tools and the arrows represent the input/output to/from each tool. ACL – anterior cruciate ligament; EMG: electromyography; GRF: ground reaction forces; MTU: muscle tendon unit.

This modified generic musculoskeletal model was then linearly scaled to match participant gross dimensions [13] using prominent bony landmarks and hip joint centres. Following linear scaling, muscle and tendon operating ranges are not necessarily preserved, so we optimized each MTU actuator’s tendon slack and optimal fibre lengths to preserve the dimensionless force-length operating curves [24]. The final step of preparing the musculoskeletal model for each participant involved updating each muscle’s maximum isometric strength as described in [25]. Then, OpenSim inverse kinematics, inverse dynamics, and muscle analysis tools were used to determine model motion, joint forces and moments, and MTU kinematics, respectively [22].

With the EMG data were conditioned and external biomechanics modelled, we used the Calibrated EMG-informed Neuromusculoskeletal Modelling (CEINMS) Toolbox [26] to estimate lower limb muscle forces. First, we used CEINMS in calibration mode for each participant using one arbitrarily selected drop-land-lateral jump and running trial. Calibration is an optimization process, described in detail [26], whereby each participant’s activation and MTU actuator parameters are adjusted, within physiological bounds, to minimize error between their joint moments calculated from inverse dynamics and those from CEINMS. We calibrated for the sagittal plane moments about the hip, knee, and ankle of the dominant (i.e., EMG instrumented) limb. Once calibrated, we used CEINMS in EMG-assisted mode [26, 27] to estimate forces for lower limb muscles in our model, as well as tibiofemoral contact forces, for all drop-land-lateral jump trials.. In EMG-assisted mode, we used the experimentally collected EMG (i.e., from the abovementioned eight lower limb muscles) as input and a novel neural control solution minimally adjusted experimentally collected EMG and synthesised excitations for the remaining 32 MTU from the lower limb. The adjusted and synthesized excitations were achieved by minimising the terms of the following objective function:

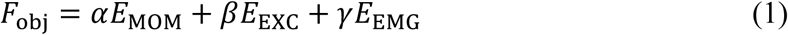

where *E*_MOM_ is the sum of the squared differences between predicted and experimental joint moments, *E*_EXC_ is the sum of squared excitations for all MTU, *E*_EMG_ is the sum of the differences between adjusted EMG excitations and recorded EMG excitations, and *α, β*, and *γ* are positive weighting coefficients [27]. We assigned value 1 to both *α* and *β* weightings and optimised *γ* weighting to minimise errors of adjusted muscle excitations and predicted joint moments.

The combined use of OpenSim and CEINMS modelling predicted lower limb muscle, intersegmental, and tibiofemoral contact forces, which were then used to estimate the ACL force via a previously validated ACL force model [13]. The ACL force model calculates the total ACL force by accounting for the planar contributions and the interaction between them:

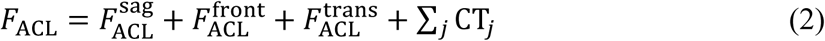

Where 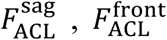 and 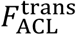 are ACL force in sagittal, frontal and transverse planes, respectively and CT_*j*_ (for *j* = SF, ST, FT) represent ACL force interactions in the sagittal-frontal (SF), sagittal-transverse (ST), and frontal-transvers (FT) planes [13].

The 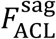 is calculated as a function of the net forces taken up by the knee ligaments 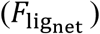 and knee flexion angle. The 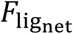 is derived from (1) knee spanning muscle forces multiplied by their line of action in anterior-posterior direction (projected onto the tibia for femur attached knee spanning MTU actuators), (2) intersegmental force in anterior-posterior direction calculated by inverse dynamics, and (3) the component of the tibiofemoral contact forces acting in anterior-posterior direction due to the posteriorly sloped tibia. The average literature values of 7.5° and 5.2° for medial and lateral tibial slope for females were used as described in [13]). Thus, 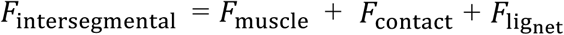 (Figure 2). The ACL force in frontal plane, 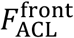, was calculated as a function of net varus/valgus moments taken up by the knee ligaments 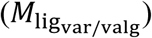 and knee flexion angle. 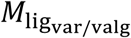 is derived from the muscle moments (calculated as muscle forces multiplied by their moment arms in frontal plane), *M*_muscle_, and intersegmental loads (knee adduction/abduction moments calculated via inverse dynamics) *M*_intersegmental_. Thus, 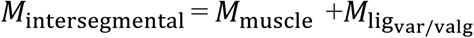 (Figure S1). Similarly, the ACL force in transverse plane 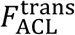 calculated as a function of net internal/external rotation moments taken up by the knee ligaments 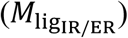 and knee flexion angle. The 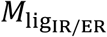 is derived from the muscle moments (calculated as muscle forces multiplied by their moment arms in transverse plane) *M*_muscle_, and intersegmental loads (knee adduction/abduction moments calculated via inverse dynamics) *M*_intersegmental_. Thus, 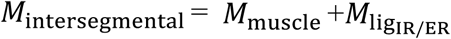 (Figure S2).

**Figure 2.**
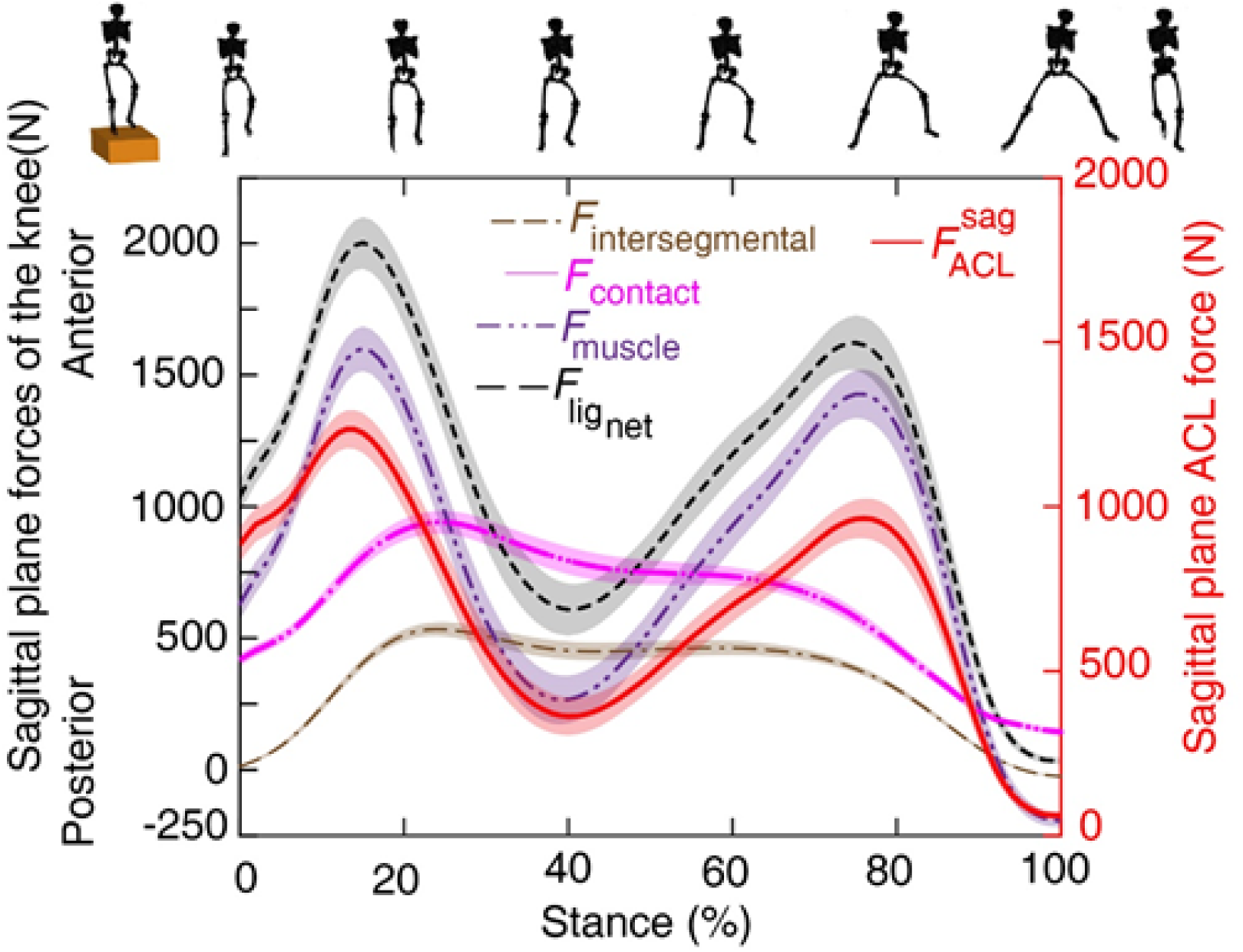
Sagittal plane knee loading (left vertical axis) and ACL force (right vertical axis) during the stance phase. Left axis: muscle forces due to their direct line of action (*F*_muscle_), tibiofemoral contact force (*F*_contact_), intersegmental forces (*F*_intersegmental_), and ligament force 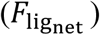 resisted by knee ligaments. 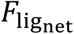 is calculated as 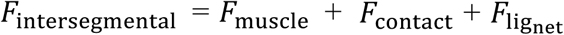. The sign of the calculated 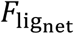 is negated as this force is resisted by the ligaments. Right axis: Force transferred to ACL through sagittal plane 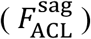. Shaded regions show standard error of the mean. Direction of sagittal plane knee load: anteriorly directed (+) and posteriorly directed (-). Silhouettes represent one participant during stance phase of the task. First and last silhouettes are before and after the stance phase, respectively, shown for clarity. ACL: anterior cruciate ligament. lig: ligament.

The abovementioned biomechanical analyses were performed across the stance phase of the drop-land-lateral jump task and normalised to 101 data points. We defined the stance phase of the drop-land-lateral jump task from initial to final foot-ground contact on the landing leg. For each participant, we analysed three trials of drop-land-lateral jump, which were then averaged to produce a single curve for the total ACL force and uni-planar contributions to this total ACL force across the stance phase. From these curves, we extracted values corresponding to the two peaks in total ACL force (raw (N) and normalized to bodyweight (BW)) and the relative contributions (% of total ACL force, reported as mean ± standard deviation) through each plane of motion. Further, at the time point corresponding to the first peak in ACL force, we grouped muscles based on their native function at the knee joint (i.e., extensor or flexors) (Table 1) and their effect on the ACL (i.e., elevating or lowering ACL force through the sagittal plane mechanism) for subsequent statistical analysis. We focused on the first peak of ACL force, since video analysis of the ACL rupture events in various sports identify the rupture to happen shortly after foot-ground (i.e., in the first 50 ms after foot-ground contact) [28].

**Table 1.**
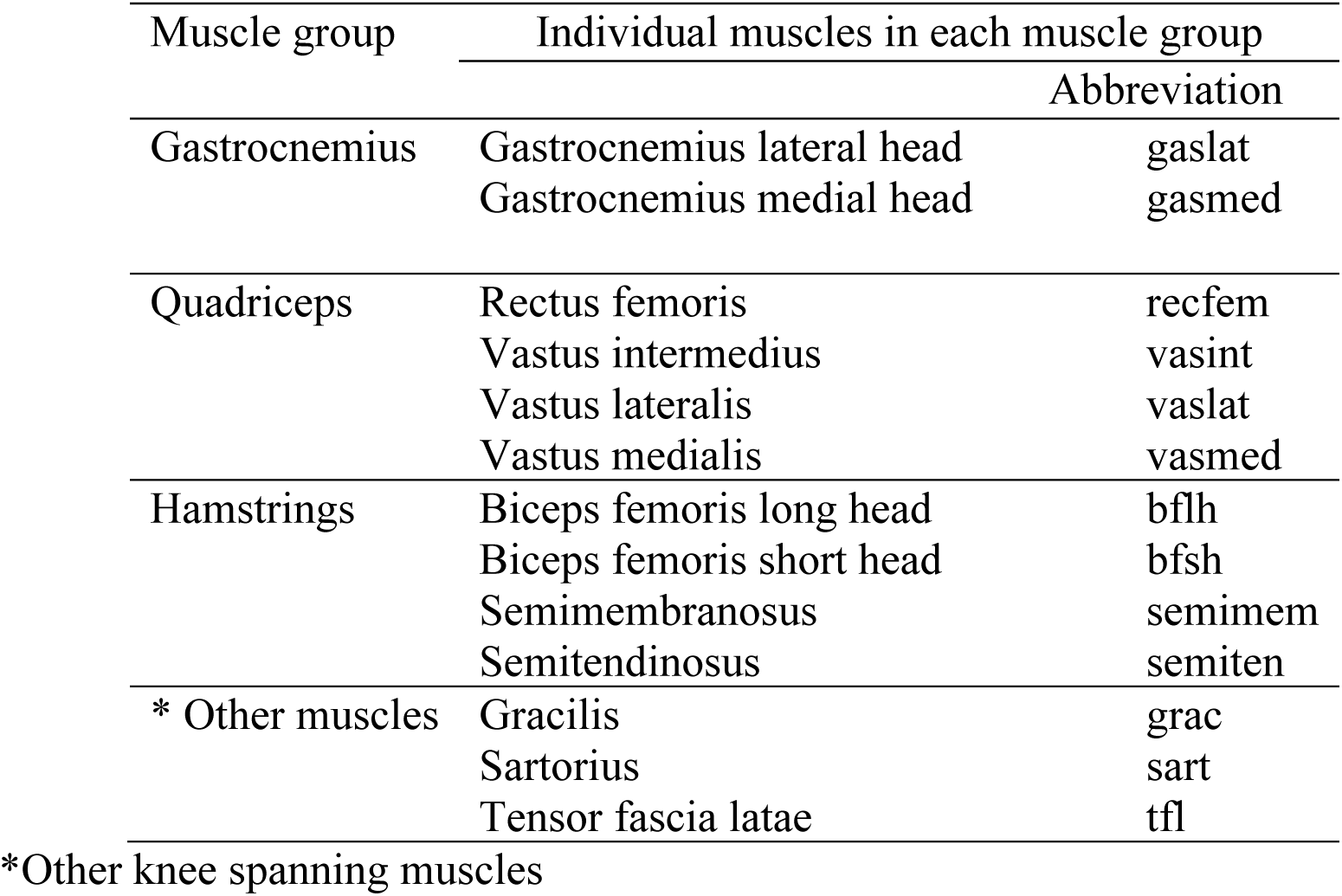
The muscle groups used for statistical comparisons.

Depending on normality of data, parametric or non-parametric one-way ANOVAs and t-tests were used to determine the differences between the relative contributions of individual muscles and muscle groups to the anteriorly directed force on the tibia caused by knee spanning muscles (*F*_muscle_) assessed at the time point of the first peak in ACL force. Statistical significance was set at p<0.05, with all analyses performed in SPSS v25 (IBM, Armonk, NY).

## Results

During drop-land-lateral jump, the majority of ACL force was delivered via the sagittal plane, followed by relatively small contributions via the frontal and transverse planes (Figure 3). We identified two peaks in the ACL force that occurred at ∼13% and 75% of stance, with the first peak in ACL force observed at 62.9±17.3 ms, shortly following initial foot-ground contact. At the first peak in ACL force, >94% of ACL force was generated through the sagittal plane and contributions from frontal and transverse planes were below 8.3% (Table 2). Note, the sagittal, frontal, and transverse planes do not sum to 100% of ACL force, as there are interaction terms specified in our model.

**Table 2.**
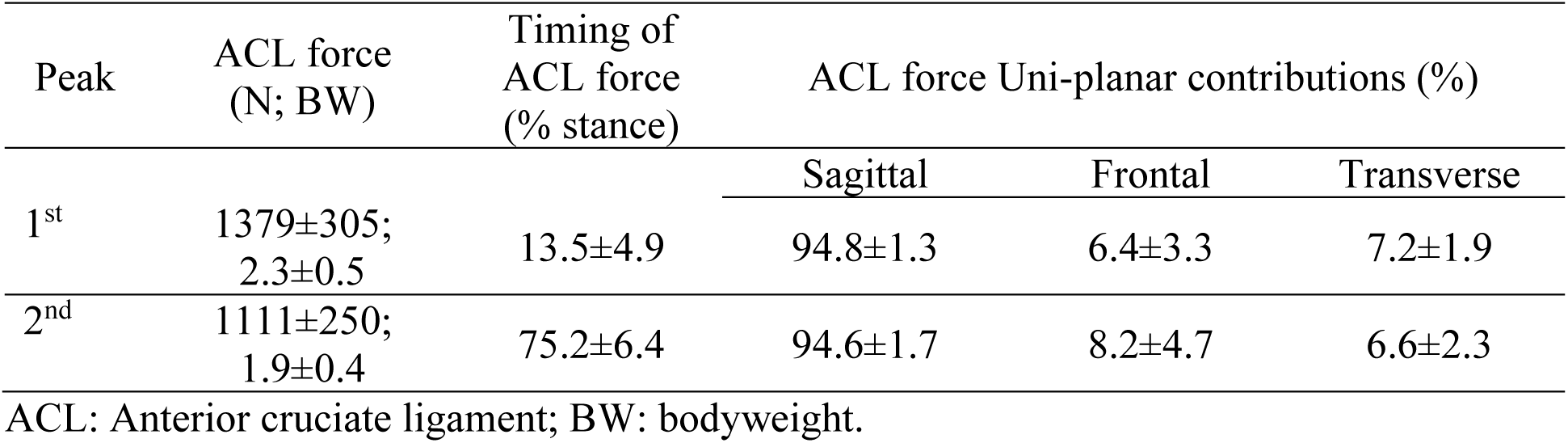
Loading parameters at the two peak points of ACL force during drop-land-lateral jump for twenty-four participants (mean ± SD).

**Figure 3.**
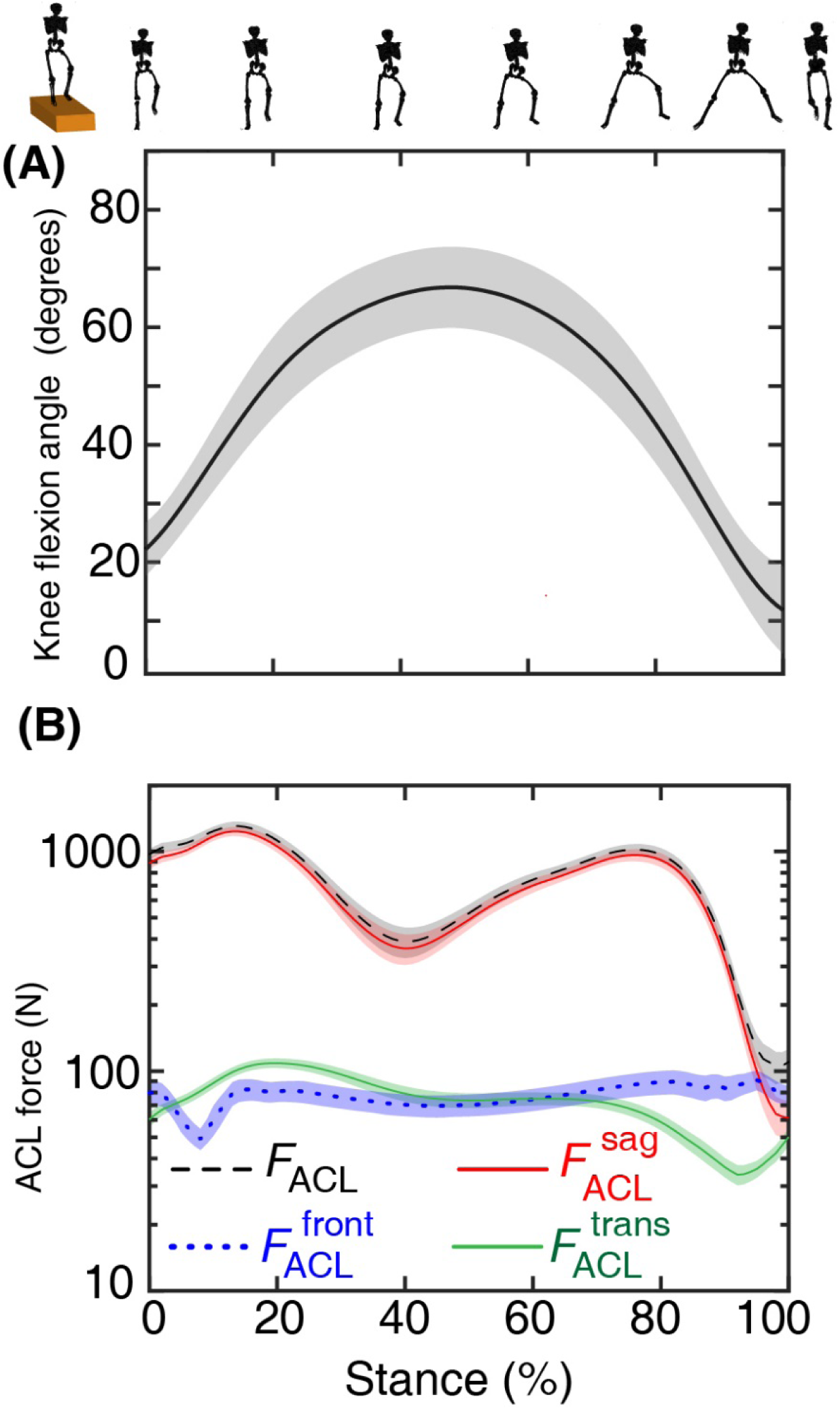
(A) Knee flexion angle and (B) ACL force across stance phase. The ACL forces developed through sagittal, 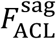, frontal, 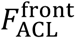, and transverse, 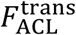, planes contribute to total ACL force *F*_ACL_. Shaded regions show standard error of the mean. Silhouettes atop subplot A represent one participant during stance phase of the task. First and last silhouettes represent posture before and after stance phase, respectively, shown to clarify the drop-land-lateral jump task performed.

In the sagittal plane, muscles were the primary contributors to the ACL force through their direct anterior-posterior lines of action, which drew the tibia anteriorly, and by compressive contact forces applied to a posteriorly sloped tibia. These forces were partially resisted by the knee ligaments (i.e., ACL, posterior cruciate ligament, medial and lateral collateral ligaments) as the net intersegmental force applied to the knee was anteriorly directed but were considerably smaller in magnitude compared to the muscle and contact forces (Figure 2). In the frontal plane, muscle generated knee valgus moment which was. partially resisted by the knee ligaments, as net intersegmental moment applied to the knee was valgus but in smaller magnitude (Figure S1). In the transverse plane, muscle generated external rotation moments about the knee that was mainly resisted by the knee ligaments as the intersegmental loads generated negligible magnitude internal rotation moments about the knee (Figure S2).

We explored the relative contributions of muscle groups (Figure 4A) and individual muscles (Figure 4B) to the anteriorly directed net muscle force (*F*_muscle_) at the first peak of ACL force. Analysis of the contributions of muscle groups revealed the gastrocnemii and quadriceps are the primary groups causing the anteriorly directed tibia force at the first peak in ACL force, providing 67.3±14.4% and 68.2±14.4% relative contributions, respectively. These contributions were not significantly different to each other (p>0.05) but were significantly larger than the other muscle groups assessed. The hamstring muscle group was the main contributor to posteriorly directed force, making -30.8 ±17% contribution. This was significantly larger (ignoring the sign convention) (p<0.05) compared to contribution from other knee spanning muscle groups (−4.7±4.6%). Figure 4B displays anteriorly and posteriorly directed forces applied to the tibia from the muscle groups at the time of peak ACL force.

**Figure 4.**
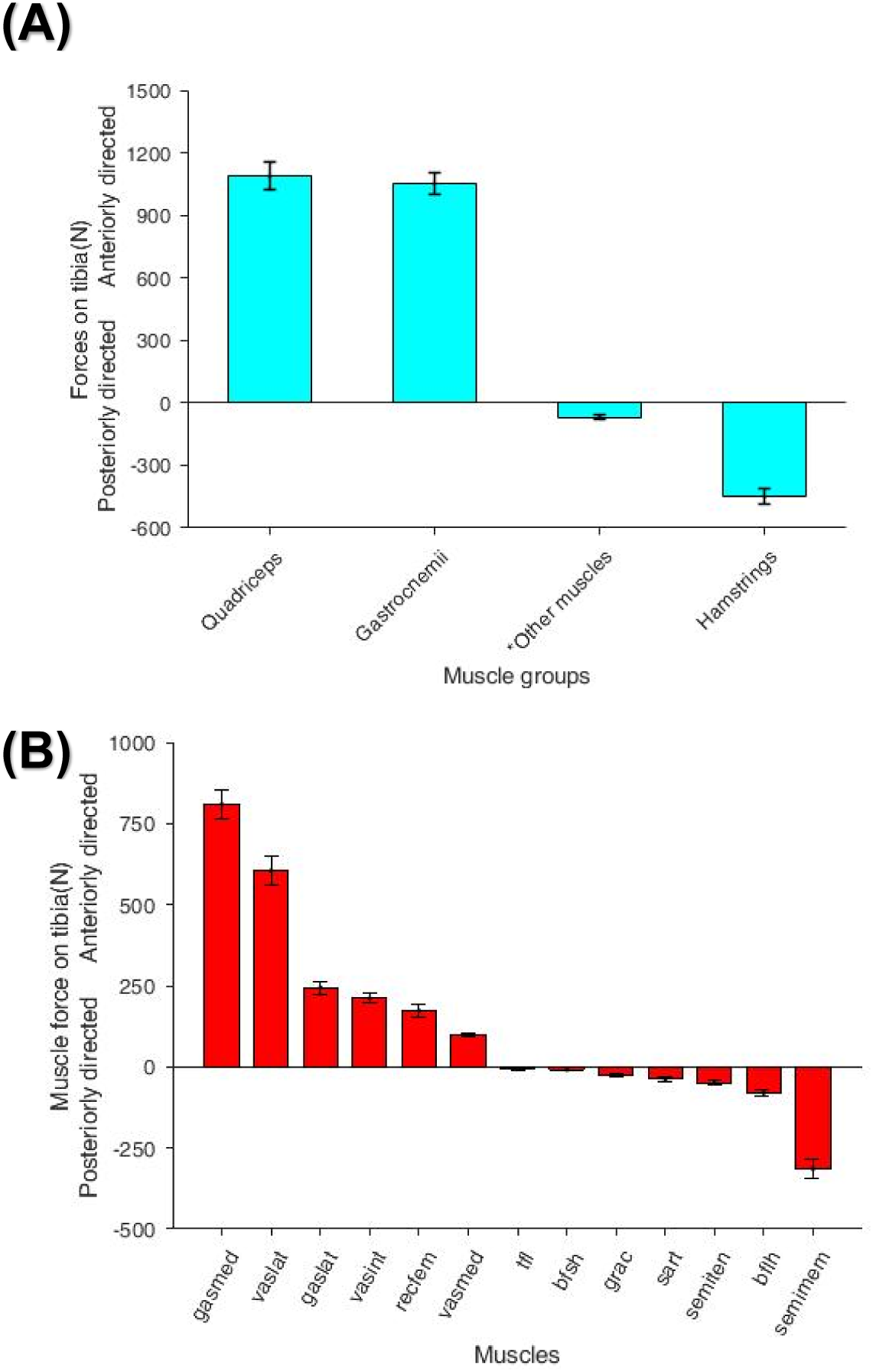
(A) Grouped and (B) individual contributions from muscle to the tibia applied through sagittal plane at the first peak of ACL force. Direction of muscle forces: anteriorly directed (+) and posteriorly directed (-). Error bars represent standard error of the mean. gasmed: Gastrocnemius medial head; vaslat: Vastus lateralis; gaslat: Gastronemius lateral head; vasint: Vastus intermedius; recfem: Resctus femoris; vasmed: Vastus medialis; bfsh: Biceps femoris short head; tfl: Tensor fasciae latae; grac: Gracilis; sart: Sartorius; semiten: Semitendinosus; bflh: Biceps femoris long head; semimem: Semimembranosus. *other knee spanning muscles (i.e., tfl, grac, and sart).

Of the individual muscle contributions, gastrocnemius medialis and vastus lateralis made the highest relative contribution accounting of anteriorly directed net muscle force at the first peak of ACL force for 51.5±12.45% and 38±11.1%, respectively. Their contributions were significantly different to each other and to all other muscles loading the ACL (p<0.05). Each of the gastrocnemius lateralis, vastus intermedius, and rectus femoris muscles made similar contributions: 15.7±6.2%, 13.5±5.2%, and 10.2±5.2%, respectively, which were not significantly different to each other (p>0.05). Vastus medialis muscles made 6.4±2% relative contribution to the anteriorly directed net muscle force which was significantly lower compared to all other muscles loading the ACL (p<0.05).

The semimembranosus was the main muscle generating posteriorly directed force to the tibia, making -21.9±14.3% relative contribution to the anteriorly directed net muscle force (Figure 4A). The negative sign signifies a posteriorly direct force (i.e., unloading the ACL). This contribution was significantly higher compared to all the other unloading contributions (p<0.05). The rest of the unloading contributions from each of the other knee spanning muscles were less than -5.5% (ignoring the sign convention) and not significantly different to each other (p<0.05).

## Discussion

The aim of this study was to determine ACL loading during a standardized drop-land-lateral jump task using a validated model that respects subject-specific muscle activation patterns, and to evaluate the contribution of knee spanning muscles, intersegmental loads, and tibiofemoral contact forces to the ACL force. To our knowledge, this is the first study to explore the ACL loading mechanism with respect to contributions from individual knee-spanning muscles using a validated multiplanar ACL force model. Results support our first hypothesis that the primary mechanism of ACL loading during the assessed task was through the sagittal plane. The gastrocnemii and quadriceps groups made the greatest relative contribution to anteriorly directed force on the tibia, thus loading the ACL, while the hamstring muscle group were the main ACL supporters applying posteriorly directed force to the tibia. These findings suggest that, in theory, by augmenting hamstring activation, ACL force could potentially be lowered. However, the effects of muscle forces on ACL loads are complex and non-linear, therefore further research using real-time ACL force for biofeedback would enable personalized strategies for ACL force reduction and may eventually reduce ACL injury.

Our model indicates ACL loading during the drop-land-lateral jump task was driven primarily through the sagittal plane, confirming our first hypotheses. This finding is consistent with modelling studies of landing tasks [13, 18], but diverges from studies of ACL failure in cadaveric models [6, 19, 29]. What reconciles the discrepant results from modelling live humans and cadaveric experiments are the vastly different knee kinematics therein. In recent cadaveric studies [19, 29], the femur specimen is mounted on an instrumented rig, multiplanar knee loads (including artificial muscle loading) are applied to the shank, and an axial compressive impact load superimposed on the shank, resulting in considerable knee motion (e.g., 27° valgus and 38° tibial rotation). Consequently, peak ACL strains were much higher than those reported for conditions where the knee remained stable and, importantly, where minimal non-sagittal loading occurs such as during standardized drop-land or jump tasks used in the present study.

The ACL experienced continuous (i.e., no instances of zero force), smooth (i.e., no rapid non-physiological fluctuations), and substantial (i.e., magnitude greater than bodyweight) force across the stance phase of this standardized drop-land-lateral jump task. The ACL force magnitude oscillated, displaying two local peaks separated by a distinct trough (Figure 3B), with a global maximum of ∼1380 N or ∼2.3 bodyweights. To compare our modelled ACL forces with direct measurement of ACL strains, we assumed an average ACL cross-sectional area (CSA) of 65 mm^2^ and Young’s modulus (*E)* of 113 MPa [30], and converted strain to force by ACL Force = (CSA ×*E* ×Strain). Accordingly, our modelled peak ACL force is higher compared to the ∼400 N that generate 5.47% peak ACL strain during a rapid deceleration task measured via strain gauges [7] and the ∼440 N generating 7% peak stain during a single leg-jump. Likewise, we model larger forces than the ∼880 N that caused 12% peak strain as measured by magnetic resonance imaging (MRI) and high speed biplanar radiography [8, 9] during a bilateral jump landing The lower strains reported by these previous *in vivo* studies might be due to the motor tasks examined. In our study, individuals dropped from a considerable height (30% of leg length, or ∼0.3 m) onto one leg, followed immediately by a nested task in the form of a lateral bound. In the previous studies, the tasks were a single leg vertical jump, single leg forward jump, or double leg jump-landing from a box [7-9], which have been less demanding compared to the task performed in this study. Moreover, there are known issues surrounding measuring the “unloaded” ACL to establish a reference length [5] and thus reported ACL strains from direct measurement during dynamic tasks may carry error. Nonetheless, our task elicited continuous ACL forces, larger than prior direct measurements, and peaked first shortly after landing and again at push off.

Our reported peak ACL force is also higher compared to other modelling studies [16, 31]. However, these previous modelling studies focused exclusively on the sagittal plane loading mechanism of the ACL, even though *in vivo* ACL loading is demonstrably generated through multi-planar mechanisms [4] as we noted in this current study. In addition, landing tasks performed in the previous studies were single or double leg drop-jumps confined to the sagittal plane. The task performed in our study was a single leg drop-lateral jump task, with more out-of-sagittal-plane loading, with a likely need for increased muscular stabilisation [32]. Furthermore, Mokhtarzadeh *et.al* used static optimization for estimation of muscle forces and found the contribution of gastrocnemii to ACL loading was very small. We used EMG-informed neuromusculoskeletal modelling, which respects the individual’s activation patterns, and found considerable gastrocnemii activation and contribution to loading the ACL. We also well accounted for the effects of joint compression on ACL loading due to the posterior slope of the tibia. It is interesting that Mokhtarzadeh *et.al*, reported larger peak GRF, but lower peak ACL force compared to our study despite similar participant masses and initial height above the ground. The larger peak GRF, could be due to their all male participants using stiffer landing strategies [16] compared to more compliant landing in our all-female participants. Notably, Mokhtarzadeh *et.al* did not report lower limb kinematics in their study, restricting comparison of the movements generated by our respective tasks. Our results however, were similar (i.e., slightly higher) to the results of ACL force reported by Kar *et.al* during a double leg stop-jump from a box in young female participants [33]. Kar *et.al* used a knee model with a viscoelastic ligament to estimate ACL tensile force. They reported peak ACL force of 1150 N (∼2 BW) occurred shortly after landing [33]. This similarity to our results could be explained by the similar participants characteristics. Our reported ACL force was slightly higher which could be explained by more provocative task performed in our study.

In our study, the two peaks in ACL force corresponded with absorption and push-off, respectively, while the trough separating the peaks roughly aligned with peak knee flexion. The first peak was of higher magnitude, occurring shortly after foot-ground contact at ∼13% of the stance phase. A prior modelling study that reported timing of peak ACL force during a single leg jump task [16] found a similar value to our study when accounting for between-study differences in the definition of movement phases. The timing of the peak ACL force in our study (67.7±19.3 ms after initial foot-ground contact) is similar to the time to rupture measured in cadaveric experiments of multi-planar loading simulations (54±24 ms) [34]. In contrast, recent *in vivo* experiments of dynamic ACL strain [8, 9] report peak loading occurs ∼50 ms before landing (i.e., before initial foot-ground contact) during a jump and land task. Video analysis of sports-related ACL injury indicate rupture occurs well after initial foot-ground contact, which is supported by destructive cadaver experiments [19, 34]. The two recent *in vivo* experiments [8, 9] appear to have assessed tasks that have an ACL loading profile markedly different to what occurs during common ACL injury events and drop landing.

Contrary to our second hypothesis, the gastrocnemii and quadriceps made the greatest contributions to the force applied anteriorly to the tibia, thus being the primary cause of ACL loading (Figure 4). Previous studies have focused on the roles of the quadriceps and hamstrings when exploring ACL injury risk [35] or modelling ACL loading [10, 11]. The importance of the gastrocnemii may have been overlooked, with few studies investigating the role of plantar flexor muscles on ACL loading [16, 17]. This omission may be because prior studies used musculoskeletal anatomical models and literature data which considered the gastrocnemii to have lines of action parallel to the long axis of the tibia and hence had no component generating anterior-posterior forces. Although modern definitions for the path of the gastrocnemii are included in our anatomical model, the dominant role of the gastrocnemii in ACL loading in this study is likely explained by the task studied. During drop-landing, ankle spanning muscles are highly recruited to generate torque/power that is essential to task performance. At the point of peak ACL force the knee was moderately flexed (39.5 ± 10°) and, in this position, the gastrocnemii can generate substantial posteriorly directed force on the femur primarily due to their direct anterior-posterior lines-of-action and second due to compression of the posteriorly sloped tibia [17].

The hamstring muscle group was the primary group responsible for generating posteriorly directed force to the tibia, supporting our second hypothesis and consistent with previous findings [16, 36]. However, a muscle’s role as an ACL agonist or antagonist can change as a function of lower limb kinematics, as the muscle’s line of action and force generating capacity changes. At the first peak of ACL force, the knee was moderately flexed (i.e., ∼40°) and, in this position, the hamstrings can apply a posteriorly directed force to the tibia to support the ACL. However, the support to the ACL they offer (Figure 4A) is offset by their vertical line of action which caused tibiofemoral compression and consequently anterior force due to the posteriorly sloped tibia. At larger knee flexion, hamstrings may provide better restraint to the tibia as their role in tibiofemoral compression also diminishes [37]. Altogether, our results show muscle’s mechanistic effects on ACL loading is complex, depending on lines of action and force production, both of which are sensitive to lower limb posture.

Overall, ACL loading is driven by complex interplay between muscle forces, body and inertial loads, joint contact forces, and the other passive soft tissues in the knee. Based on our findings, the hamstrings are the primary ACL supporters, thus hamstring recruitment might be a viable target for reducing load on the ACL [38]. Indeed, low hamstring-to-quadriceps strength ratio has been suggested to increase risk of ACL injury [39]. However, it is not clear how hamstring recruitment during dynamic landing tasks can readily be increased and retained. Some studies have shown isometric strength or balance training is ineffective in increasing hamstring coactivity with the quadriceps [40, 41], others suggest specific strength training can increase hamstring-quadriceps coactivation and potentially be beneficial in ACL injury prevention and rehabilitation [42]. Thus, the selection, intensity, and biofeedback of exercises might be important when focusing on ACL injury prevention, as has been suggested in literature [43, 44]. In all cases, the relationship between EMG measures and muscle force production are highly tenuous, requiring sophisticated neuromusculoskeletal modelling to verify mechanical effects on the ACL.

The current study has several limitations that should be acknowledged. First, we used a scaled version of a generic musculoskeletal model to perform our modelling and simulations. Currently, the effects of increasing the level of model personalization on modelled ACL force is unclear, but presumably a better match to the individual from anatomical and neuromuscular perspectives will provide superior predictions of internal body mechanics. Second, the drop-land-lateral jump task performed in this study was planned which unlikely represents the patterns of body motion, external loading, and muscle activations present when a non-contact ACL injury occurs. Furthermore, this task failed to elicit large external valgus at the knee, which appears in many video recordings of ACL ruptures in sport. Therefore, the task may have not been major challenge for the neuromuscular and ligamentous restraints at the knee. Future research may consider more provocative tests of knee stability if they are intended to reveal those who can well stabilize the knee and those who cannot, with attention to task standardization. Despite these limitations, our study did find the time to peak ACL force consistent with video analysis of injury and destructive cadaveric testing. Importantly, our analysis does clarify how ACL forces are generated during a standardized movement task, which provides further understanding of the natural loading mechanisms of the ACL.

## Conclusion

Our results demonstrated the ACL is loaded primarily through the sagittal plane during a standardized drop-land-lateral jump task. In the sagittal plane, muscles generated substantial forces directed anteriorly to the tibia and hence ACL force through their direct lines of action. Muscle contribution to ACL force far outweighed that of the tibiofemoral contact forces. The gastrocnemii, along with the quadriceps, were the main ACL loaders, while hamstrings were the major supporters of the ACL. Results highlight the important role of gastrocnemius muscle in ACL loading, which may have previously been overlooked and may be considered more prominently in ACL injury prevention and rehabilitation programmes.

