## Supplement for "Mechanism of anterior cruciate ligament loading during dynamic motor tasks"

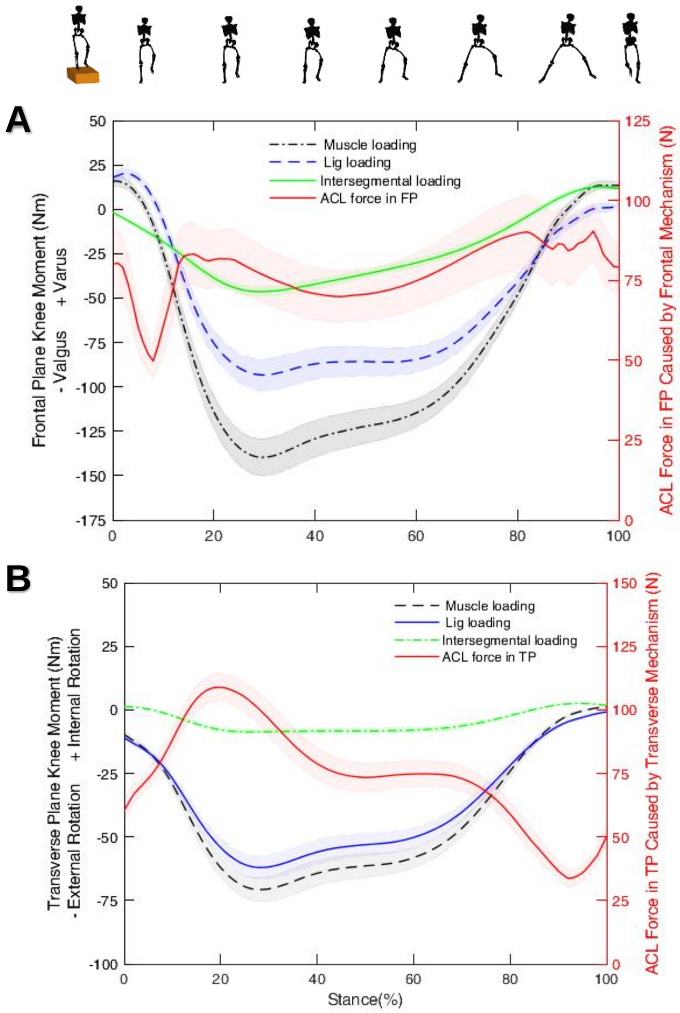


Figure S1. Uni-planar ACL forces across the stance phase of the drop-land-lateral jump task. (A) Frontal plane, and (B) transverse plane knee loadings and ACL forces. Left axis: muscle and intersegmental moments ($M_{\text{muscle}}$ and $M_{\text{intersegmental}}$), and net varus or valgus moment

$M_{\text{lig}_{\text{var/valg}}}$, and internal or external rotation moment $M_{\text{lig}_{\text{IR/ER}}}$ resisted by the knee ligaments; Right axis: ACL forces $F_{\text{ACL}}^{\text{front}}$ and $F_{\text{ACL}}^{\text{trans}}$. The shaded regions represent the standard error of the mean. Direction of frontal plane knee moment: varus (+) and valgus (-); transverse plane knee moment: internal rotation (+) and external rotation (-). Silhouettes represent one participant during the stance phase of the task. First and last Silhouettes are before and after the stance phase, respectively, shown for the clarity. ACL: anterior cruciate ligament; FP: frontal plane; SP: sagittal plane; Lig: ligament.
